# ODC1 restricts meningeal B cell age-associated-like phenotype and function in multiple sclerosis: a human and experimental study

**DOI:** 10.1101/2025.09.09.674821

**Authors:** Jonathan Zurawski, Martin R. Profant, Alara Tuncer, Jianuo Wang, Miranda Green, Ke Cao, Simon Paris, Shahamat Tauhid, Youmna Jalkh, Molly Quattrucci, Renxin Chu, Xingshan Cao, Alex Kiss, Tanuja Chitnis, Howard Weiner, Clary Clish, Rohit Bakshi, Chao Wang

**Author notes:** Joint first authors (Jonathan Zurawski, Martin R. Profant, Alara Tuncer).

## Abstract

Meningeal inflammation, as a clinical feature of multiple sclerosis (MS), is associated with worse clinical disease outcomes. In both relapsing and secondary progressive MS and the experimental autoimmune encephalomyelitis (EAE) MS model, the meninges have been found to contain ectopic lymphoid follicles enriched with B cells. The metabolic requirement of meningeal B cell function in MS or EAE is not well elucidated. Using 7-Tesla MRI brain scans of MS patients and leptomeningeal enhancement (LME) as a marker, we found a correlation between meningeal inflammation and metabolites of the arginine/polyamine pathway, a finding recapitulated in the EAE model. Ornithine Decarboxylase (ODC1), the rate limiting enzyme for polyamine biosynthesis, as well as polyamine metabolism was diminished in the dura meningeal B cells from mice with MOG_35-55_ induced EAE mice as compared to naïve controls. Pharmacological inhibition of ODC1 restricted meningeal T cells but promoted meningeal B cell proliferation. B cell-specific deletion of ODC1 resulted in expansion of B cells with age-associated B cell-like phenotype (CD11c^+^CD21/35^-^CD23^-^IgD^-^) and exacerbated disease in the MOG_1-125_ EAE model. Together, these findings demonstrate a divergent role of polyamines in regulating B and T cell responses in the meninges during autoimmunity.

**Significance Statement:** This study identified the polyamine pathway to be associated with meningeal inflammation in multiple sclerosis, a clinical phenotype associated with worse disease outcomes and without targeted therapy. Using a mouse experimental model, we found that ODC1, the rate limiting enzyme of the polyamine biosynthesis pathway, was suppressed in meningeal B cells, restricted the development of age-associated B cells in the meninges and limited disease severity. This study elucidated a metabolic pathway regulating meningeal B cell function, informing its therapeutic applications in autoimmune diseases.

## INTRODUCTION

Meningeal inflammation is a hallmark of multiple sclerosis (MS), detectable early in the disease course, and more frequently in patients with progressive MS by histology.^1–4^ This finding is further supported by neuroimaging data acquired from post-contrast brain MRI, which enables visualization of leptomeningeal enhancement (LME), a proposed marker of meningeal inflammation.^5–11^ Animal models of MS have shown that CNS penetrant therapy targeting myeloid cells and B-cells limits meningeal and adjacent cortical inflammation.^12^ In people with MS, while meningeal inflammation is hypothesized to contribute to disease progression, there are no therapies available to target this process directly.

Meningeal inflammation is characterized by ectopic lymphoid follicles (eLFs) enriched with germinal center-like structures, and meningeal B cell follicles have been demonstrated to be a feature of the pathology of secondary progressive MS.^13^ In the experimental autoimmune encephalomyelitis (EAE) model of MS, meningeal eLFs are enriched with B cells, effector T cells, regulatory T cells and granulocytes,^14^ supported by a fibroblastic reticular cell network.^15^ Meningeal B cells are enriched with a unique repertoire and show evidence of affinity maturation in the EAE model.^16^ Previous studies show that CD4^+^ T helper 17 cells (Th17) can drive the formation of eLFs in EAE.^17^ This process depends on the expression of Bcl6 in Th17 cells,^18^ and is promoted by lymphotoxin αβ signaling between Th17 and meningeal stromal cells.^15^ Myelin-reactive meningeal B cells can also produce IL-23, further promoting the expansion of pathogenic Th17 cells.^19^ Indeed, the antigen-specific interaction between meningeal B cells and Th17 cells can result in long-lasting reactivation of autoreactive T cells and B cell differentiation.^20^ The cooperation between meningeal B and T cells is thought to propagate CNS inflammation.

Cellular metabolism is a critical regulator of both B and Th17 cell differentiation and function.^21,22^ We and others have previously shown that Th17 cell differentiation and proinflammatory function are dependent on metabolic regulators such as HIF1a,^23^ glucose metabolism via PDHK1,^24^ GLUT3,^25^ and PGAM1;^26^ fatty acid biosynthesis via CD5L^27^ or ACC1;^28^ amino acid metabolism via glutaminase (GLS) ^29^ and polyamine metabolism via Ornithine Decarboxylase 1 (ODC1) and diamine acetyltransferase (SAT1).^30^ This diverse metabolic network is likely driven by a complex interplay between cytokines such as IL-6, IL-1b, TGFb, and IL-23 together with TCR signaling. Similarly, germinal center (GC) B cells are supported by metabolic reprogramming induced by the IL-4-Bcl6 axis.^31^ CD40 signaling further promotes the expression of MTHFD2 that drives one-carbon metabolism and maintains the redox state necessary for GC B cell survival and proliferation.^32^ Glycolysis-dependent 3-phophoglycerate production fuels the serine biosynthesis pathway via the rate limiting enzyme, PHGDH, required for GC B cell activation and affinity maturation.^33^ In contrast to clonally expanding T cells or pre-GC B cells, ex vivo GC B cells show dependence on fatty acid oxidation instead of glycolysis for ATP production. ^34,35^ The metabolic requirement of meningeal B cells and mechanisms promoting their activation remains unclear.

In this study, we found that plasma levels of arginine-polyamine metabolites positively correlated with meningeal inflammation and inversely correlated with normalized grey matter volume as measured by 7-Tesla (7T) brain MRI in people with MS. This finding was recapitulated in the MOG_35-55_ EAE model, where the numbers of meningeal GC-like B and effector T cells significantly correlated with hippocampal polyamine levels and hippocampal volume. Consequently, we further investigated the function of the polyamine pathway in regulating meningeal inflammation. Surprisingly, while inhibition of the rate limiting enzyme of the polyamine biosynthesis (ODC1) limited the number of meningeal effector T cells, it expanded dural GC-like B cells. We found that in contrast to Th17 cells, meningeal B cells suppressed polyamine metabolism upon activation. ODC1 perturbation promoted meningeal B cell proliferation in vitro and an expansion of age-associated-like B cells in dural meninges in vivo. Consistently, B-cell intrinsic deletion of ODC1 resulted in worse EAE using the MOG_1-125_ protein immunization model, pointing to a suppressive role of ODC1 in meningeal B cell immunity.

## RESULTS

### Plasma metabolic profile is associated with brain grey matter volumes and meningeal inflammation of MS

To evaluate any associations of blood metabolic markers to meningeal inflammation in MS, we acquired plasma samples from a cohort of 68 MS patients from the CLIMB study.^36^ Clinical characteristics are shown in **Table 1**.

**Table 1.** Clinical-demographic characteristics and MRI variables of an MS subject cohort with 7T MRI and blood obtained.

Data are mean ± standard deviation (range) unless otherwise indicated; DD = disease duration in years from symptom onset; RRMS = relapsing-remitting multiple sclerosis; SPMS = secondary progressive MS; PPMS = primary progressive MS; CIS = clinically isolated syndrome; EDSS = Expanded Disability Status Scale.
|  | Untreated (n=23) | Treated (n=45) |
| --- | --- | --- |
| **Age (years)** | 51.4 ± 12.4 | 52.5 ± 11.7 |
| **Female (%)** | 78% | 77% |
| **DD (years)** | 16.2 ± 11.2 | 16.3 ± 10.8 |
| **RRMS, n(%)** | 16 (70%) | 28 (62%) |
| **SPMS, n(%)** | 3 (13%) | 10 (22%) |
| **PPMS, n(%)** | 2 (9%) | 6 (13%) |
| **CIS, n(%)** | 2 (9%) | 1 (2%) |
| **EDSS** | 3.1 ± 2.3 | 3.5 ± 2.4 |

Leptomeningeal enhancement (LME) prevalence was 67% (n=44) for the overall cohort (**Figure 1 and Table 2**). Higher LME prevalence was seen in progressive vs. RRMS (80% vs. 64%) subjects and in anti-CD20-treated vs. untreated (73% vs. 55%) subjects. LME foci number correlated positively with EDSS (r=0.47) and inversely with total deep gray matter (DGM=caudate+putamen+globus pallidus+thalamus) volume (r=-0.23), both p<0.05.

**Figure 1.**
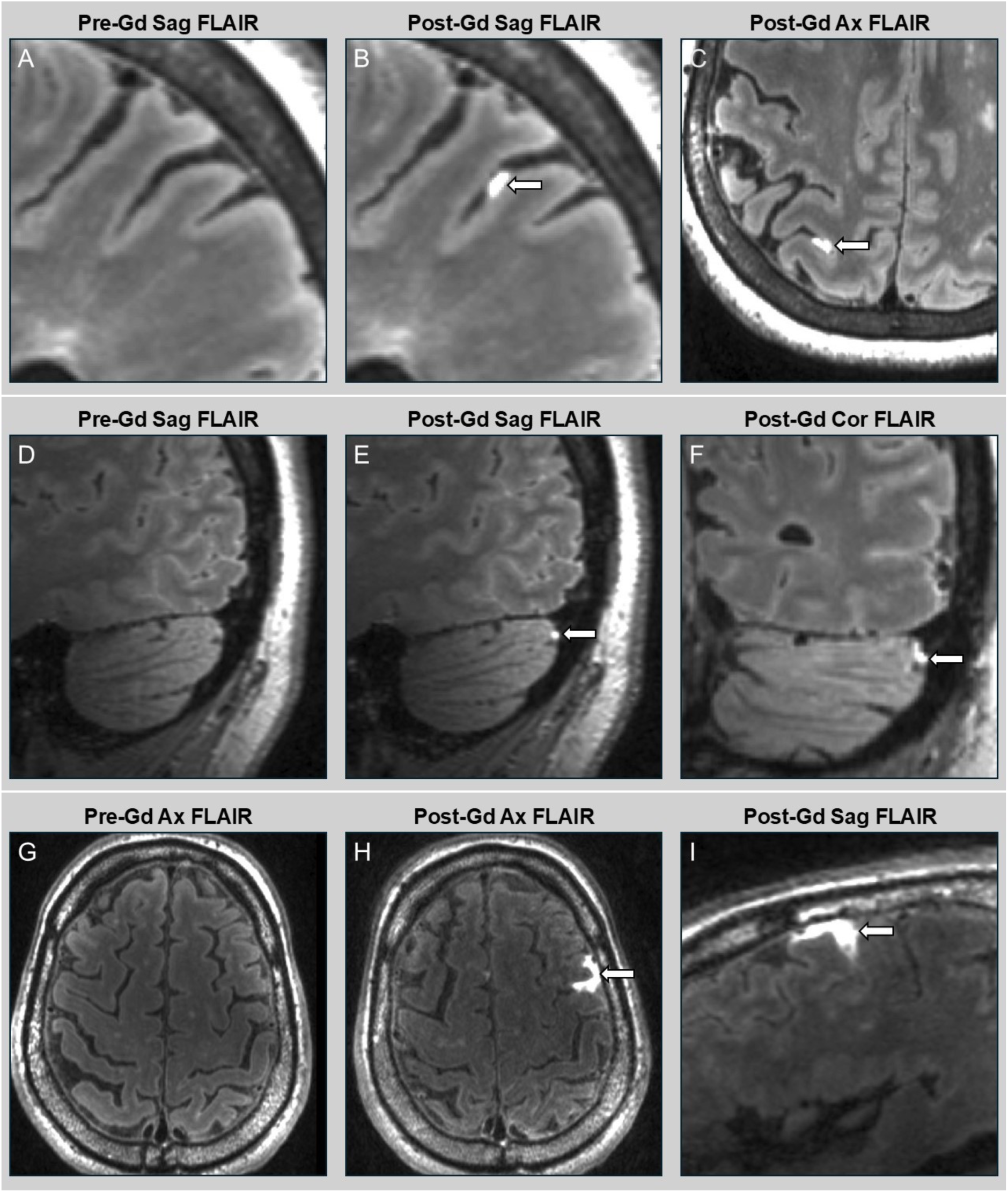
Representative Leptomeningeal Enhancement in MS Subjects. Representative leptomeningeal enhancement (LME) from the 7T Siemens Terra Magnetom at Brigham & Women’s Hospital, three MS patients treated with anti-B cell therapy are shown. Top panel (**A-C**): A 66-year-old woman with secondary progressive MS on rituximab with a right parieto-occipital nodular focus of LME on post-contrast sagittal (B) and axial (C) FLAIR sequences (white arrows indicate LME focus). Pre-contrast sagittal FLAIR sequence (A) shows no contrast enhancement in the corresponding anatomical region. Middle panel: (**D-F**) A 51-year-old woman with relapsing MS treated with ocrelizumab with a nodular focus of LME (arrow) in the cerebellum evident on post-contrast sagittal (E) and coronal (F) FLAIR sequences. Lower panel: (**G-I**) A 74-year-old man with secondary progressive MS treated with ocrelizumab with a large spreading focus of LME (arrow) in the left temporal lobe seen on (**H**) axial and (**I**) sagittal FLAIR sequences. Sag = sagittal; Gad = gadolinium; Ax = axial; Cor = coronal.

**Table 2.** MRI Variables.

Data are mean ± standard deviation (range) unless otherwise indicated; LME = leptomeningeal enhancement; DGM = deep gray matter; T2LV = White Matter T2 lesion volume; BPV = brain parenchymal volume.
|  | Untreated | Treated |
| --- | --- | --- |
| **LME+, n (%)** | 12 (55%) | 32 (73%) |
| **LME foci number** | 1.4 ± 1.5 (0-4) | 2.2 ± 2.2 (0-9) |
| **T2LV (ml)** | 4.5 ± 4.6 | 6.0 ± 8.5 |
| **BPV (ml)** | 1492 ± 123 | 1500 ± 95 |
| **Total DGM volume (ml)** | 44.1 ± 4.9 | 44.2 ± 4.0 |
| **Thalamus volume (ml)** | 18.9 ± 2.4 | 18.9 ± 1.8 |
| **Caudate volume (ml)** | 9.2 ± 1.2 | 9.2 ± 1.0 |
| **Putamen volume (ml)** | 11.9 ± 1.5 | 11.9 ± 1.2 |
| **Globus Pallidus volume (ml)** | 4.3 ± 0.4 | 4.3 ± 0.6 |
| **Hippocampus volume (ml)** | 8.9 ± 1.3 | 9.2 ± 1.2 |

We performed targeted metabolomics of the plasma and obtained quantification data for 377 metabolites (**Methods**). To assess the associations between metabolites and MRI imaging data, including LME foci number, T2 hyperintense white matter lesion volume (T2LV), DGM, thalamus, caudate, putamen, globus pallidus and hippocampus volumes, we constructed an unbiased sPLS model that optimized both the number of metabolite (X) and imaging variables (Y) features to reduce dimensionality while maximizing covariation between inputs and outcomes. Of the metabolites quantified, those with 30% missing values were excluded and a total of 357 metabolites were included in the final analysis (**Methods**). The ensemble model, which chooses iterations with the lowest error rate in each round of cross-validation, selected 107/357 and 258/357 metabolite variables in the first and second component, respectively, with an average of 36 and 56 variables selected by each independent model (listed in Table S1). In the X-variate space, three fatty acyl carnitines (3-methylglutaroyl-carnitine [CAR(5:0-DC)], acylcarnitine(C9:0) [CAR(9:0)] and acylcarnitine(C20:0) [CAR(20:0)]) had the highest positive loading weights along principal component (PC) 1, while methylimidazole acetic acid, N-acetyl serine, and allantoin had the highest negative weights, suggesting that these metabolites account for the largest source of variation in imaging data across patients (**Figure 2A**). All MRI imaging variables (LME, T2LV, DGM, thalamus, caudate, putamen, globus pallidus, and hippocampus volumes) were selected across both components of the average ensemble model (**Figure 2B**). Notably, we found that markers of atrophy including regional brain volumes (DGM, hippocampus, and caudate) were selected for model building across the first two components in all 50 model iterations, while the remainder (putamen, thalamus and globus pallidus) were selected at a lower frequency, and measures of inflammation, LME foci number and T2LV, were selected for 12 and 15 models, respectively (inclusion rates summarized in Table S2). This was also reflected in average loading values, where T2LV and LME foci number had the lowest absolute overall contribution to the Y-variate space of the ensemble model (**Figure 2B**). Furthermore, multivariate decomposition of the metabolite (X) and XY variate space (all selected metabolite and imaging features) showed no systematic variation across patients with different LME foci number and T2LV (**Figure 2C**). Overall, these results indicate that peripheral metabolomics profiles are more strongly associated with specific regional brain volumes than with inflammatory markers of MS progression. They also reaffirm that different outcome measures contribute unequally to the overall model, reflecting divergent relationships with selected metabolites.

**Figure 2.**
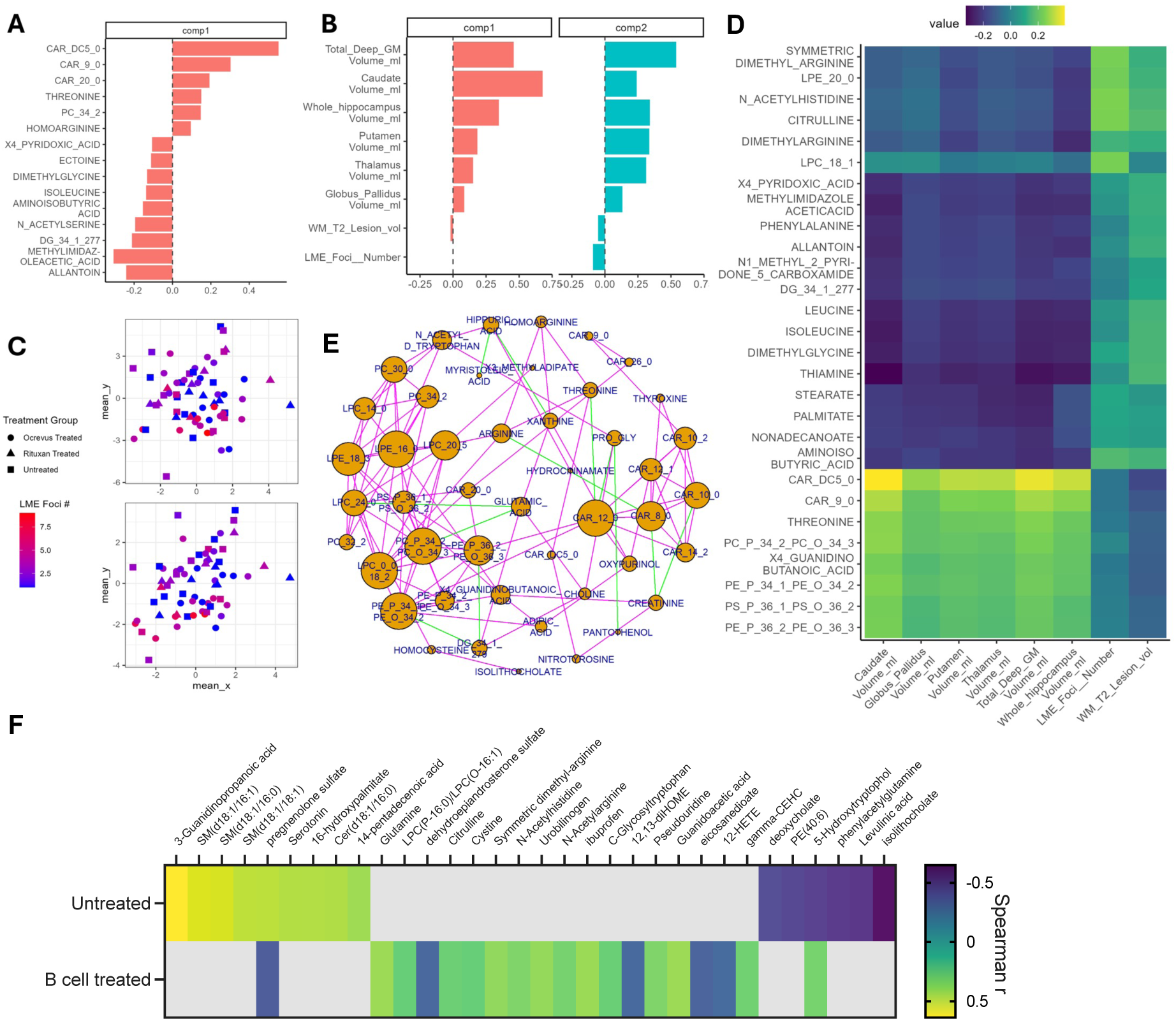
Metabolomic landscape of MS patients and its relationship to brain imaging data. **A)** top 15 average X-variate loadings along the first principal component (PC) of the ensemble sPLS model arranges by descending value. **B)** Average Y-variate loadings across the first principal component (PC) of the ensemble sPLS model. **C)** sPLS dimension reduction of the metabolite block (X, top), and X-Y variate space (metabolites and imaging, bottom) with each data point colored according to LME foci count and representing treatment groups by shape. **D)** Clustered Image Map (CIM) representing the first 2 components of the ensemble sPLS model. Each value in the heatmap (the correlation of each original feature pair) is determined by each of their correlations with the components from the integrative method. A cutoff of 0.2 was applied to pairwise metabolite-image feature associations. **E**) network plot showing the network associations between metabolites in the first principal component of the sPLS model (spearman rank correlation, R>0.3 and adjusted p-value < 0.05). Green lines represent negative correlations while magenta represents positive correlations. **F)** Spearman correlation analysis was performed between LME foci number and metabolites for either untreated or B-cell therapy treated cohorts. Only significant correlations (p < 0.05) are reported in the heatmap which visualizes *Spearman* r. Grey grids are non-significant (p > 0.05).

### Arginine derivatives correlate with meningeal inflammation most closely in MS patients treated with B cell therapy

To directly investigate the metabolites in covariation with meningeal inflammation and brain volume, a subset of model variables were further visualized via a clustered imaging map (CIM) along the first two PCs of the ensemble model (pairwise correlation cutoff value >0.3, **Figure 2D**). This approach illustrated positive correlations between regional brain volumes and a cluster of compounds including the acyl carnitines (CAR(X:X)), glycerophospholipids (PE(P-X) or PC(P-X)), threonine, and metabolites connected to the arginine-polyamine pathway including creatinine and 4-guanidinobutanoic acid. By contrast, LME foci number and T2LV were most strongly correlated with arginine derivatives (symmetric dimethyl arginine [SDMA], citrulline, dimethyl arginine) which were negatively associated with differences in regional brain volumes. Together, these results suggest that distinct metabolite clusters, even those within the same pathways such as the arginine-polyamine pathway, show divergent associations with regional brain volumes and inflammatory markers, suggesting unique roles in MS progression.

Next, to examine the associations between specific metabolites, we constructed a correlation network, filtering significant correlations (p_adj_ < 0.05) between metabolites along the first principal component of the ensemble model with a threshold of |R| > 0.3 (**Figure 2E**). We observed several positive associations (magenta lines) between glycerophospholipids (PE(P-X) or PC(P-X)), lysophospholipids (LPE(X:X) and LPC(X:X)), and fatty acyl carnitines (CAR(X:X)). Highly connected clusters of the lipid derivatives were in turn negatively associated (green lines) with metabolites such as creatinine, proline-glycine, arginine, glutamic acid, threonine and choline, indicating negative covariance between lipid and amino acid derivatives in the overall metabolomic landscape.

Finally, we analyzed untreated and B-cell therapy treated cohorts separately, focusing on metabolites correlated with LME foci numbers (**Figure 2F**). We found that meningeal inflammation correlated with a distinct set of metabolites in the two cohorts: in untreated patients, we observed correlation with mostly lipid derivatives (e.g. sphingomyelins, and ceramide); whereas in patients treated with B cell therapies derivatives of arginine and other amino acids were the major correlated metabolites including those not captured by the overall model (e.g. guanidoacetic acid and N-acetyl arginine, both derivatives of arginine). Overall, there was a stronger correlation of arginine derivatives with LME foci number in the treated cohort, suggesting a connection between these metabolites and B cell therapy.

### Brain polyamine levels increase in chronic EAE and are associated with meningeal B and T cell expansion

The association of blood arginine/polyamine metabolite levels with meningeal foci and grey matter volume in MS prompted further investigation of this pathway and its effect on meningeal immune cells in the MOG_35-55_ EAE model. To this end, we first validated the model by assessing meningeal inflammation comparing naïve, onset and chronic phase (days 0, 10-14, and 54-60) of EAE in wildtype C57BL/6 mice (**Figure 3A**). We conducted a flow cytometric analysis of dural and leptomeningeal tissue. We found a significant increase in the number of effector CD4 T cells (Teff, CD45^+^CD3^+^CD4^+^Foxp3^-^) in dural meninges at onset (day 14) of EAE in both male and female mice which returned to baseline by day 54 (**Figure 3B**, upper left panel). In contrast, in the leptomeninges, Teff cells drastically expanded only at day 54 (**Figure 3B**, upper right panel). Notably, despite the T-cell driven nature of the MOG_35-55_ model, we observed a significant accumulation of GC-like B cells (CD19^+^GL7^+^FAS^+^) in the leptomeninges but not the dura, which coincided with Teff expansion (**Figure 3B**, lower panels).

**Figure 3.**
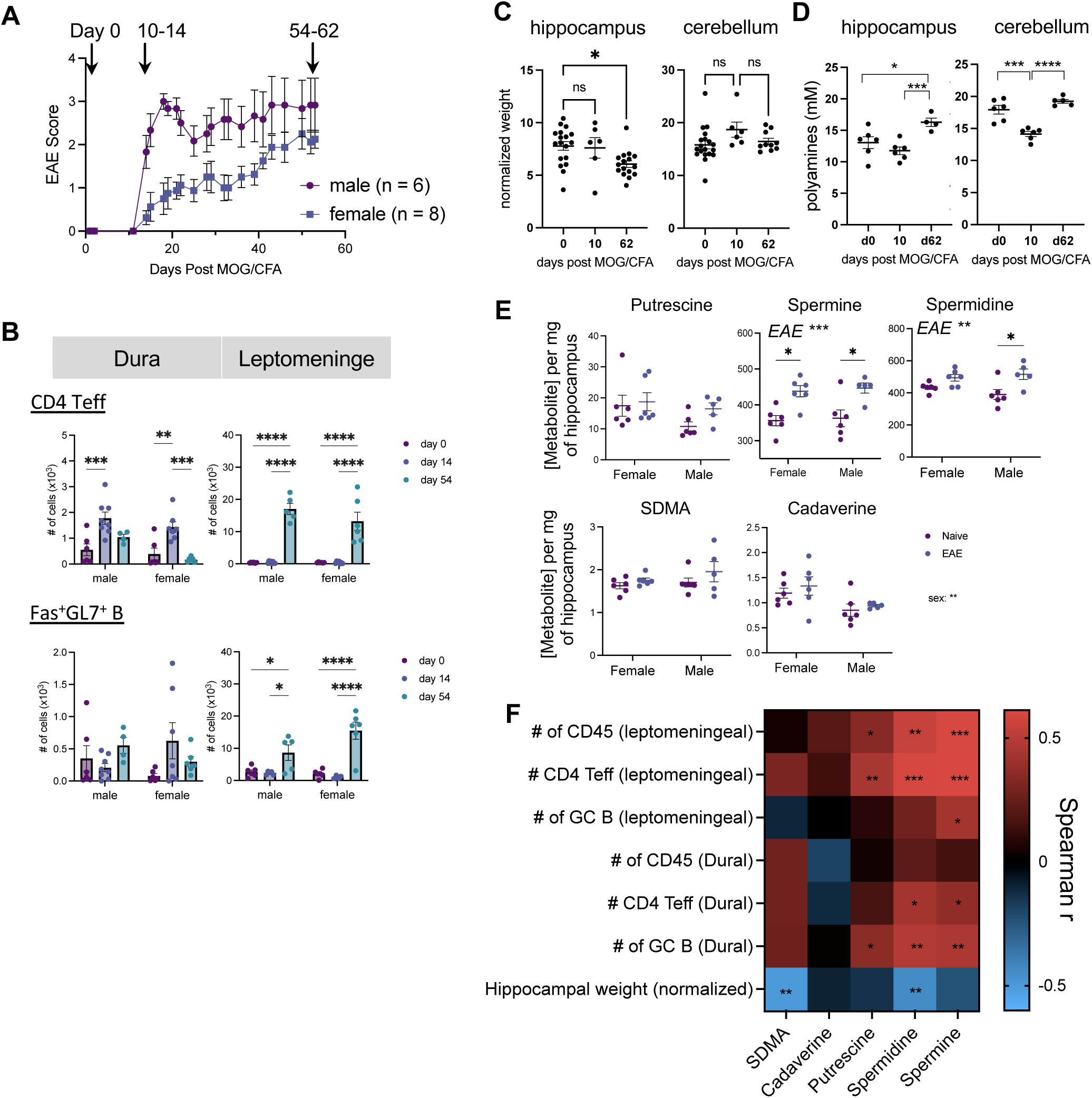
Polyamines are associated with EAE induced meningeal and hippocampal dysregulation. C57BL/6 wildtype mice of both sexes were immunized with MOG_35-55_/CFA to induce EAE. **A)** Representative EAE score where naïve (day 0), onset (day 10-14), and chronic phase (day 64-62) were indicated. **B)** Dural and leptomeningeal tissue were dissected kinetically from EAE mice. CD4 Teff (CD45^+^CD3^+^CD4^+^Foxp3^-^) and GC-like B cells (CD19^+^GL7^+^FAS^+^) were quantified using counting beads and flow cytometry. **C-D)** Hippocampus and cerebellum were dissected, weighed and normalized against whole brain weight (C). Interstitial fluid from hippocampus were quantified for total polyamine using enzymatic assay (D). **E)** Interstitial fluid from hippocampus (EAE, day 54) were analyzed by targeted metabolomics using Mass Spectrometry. **F)** Correlation matrix of metabolomics data acquired in (E) against either the number of meningeal immune cell or hippocampi weight acquired similarly to B and C. Each dot represents data from one mouse. Statistical analysis: B and E used Two-way ANOVA with Tukey’s multiple comparisons test; C and D used one-way ANOVA with Dunnett’s multiple comparisons test. * p < 0.05; ** p < 0.01; *** p < 0.001; **** p < 0.0001.

Next, we investigated changes of polyamine metabolites in the EAE brain. Given that hippocampal volume showed the strongest association with peripheral metabolic changes in brain MRI among people with MS, we focused on this region. In EAE mice, we found a loss of normalized weight of hippocampus, but not cerebellum, at the chronic phase (day 62), highlighting the hippocampus as a region particularly vulnerable to inflammation (**Figure 3C**). Using an enzymatic assay, we identified increased levels of total polyamines in the hippocampal tissue, whereas the cerebellum had a temporary decrease at disease onset that returned to baseline by day 62 of EAE (**Figure 3D**). To better resolve the types of polyamines involved, we performed targeted metabolomics of the interstitial fluid of hippocampus tissue isolated from naïve mice and mice at day 62 of EAE. We found that spermine and spermidine, but not putrescine, SDMA, or cadaverine, were increased in chronic EAE (**Figure 3E**).

Finally, we performed a correlation analysis for male and female EAE (day 62) mice comparing polyamines levels, meningeal immune cell numbers, and hippocampal volume. We found that while normalized hippocampal weight was negatively associated with the level of SDMA and spermidine, both Teff and GC-like B cells were positively correlated with all three major polyamine metabolites (putrescine, spermine, and spermidine), with some nuisance effect of whether present in leptomeninges or dura respectively (**Figure 3F**).

### Polyamine metabolism is active in naïve meningeal B cells and suppressed in EAE

To study the role of the polyamine pathway in meningeal B cells at steady state and in EAE, we performed single cell RNAseq (scRNAseq) and assessed the polyamine pathway using both gene expression and Compass, an algorithm we previously published to model metabolic flux within the scRNAseq data^30^ (**Figure 4**). UMAP visualization of 19613 dura meningeal B cells isolated from naïve mice and mice at disease onset (day 14) of MOG_35-55_ EAE revealed distinct transcriptional clusters (**Figure 4A**). Consistent with their identify, cells expressed B cell-related genes (e.g. *Ighm, Ighd, Ms4a1, CD74, H2-Aa, Rag1, and IL7r*) that allowed us to distinguish between developing B cells (clusters 1, 3, and 4) and mature B cells (clusters 0, 2, 5-8) in our dataset (**Figure 4B**).

**Figure 4.**
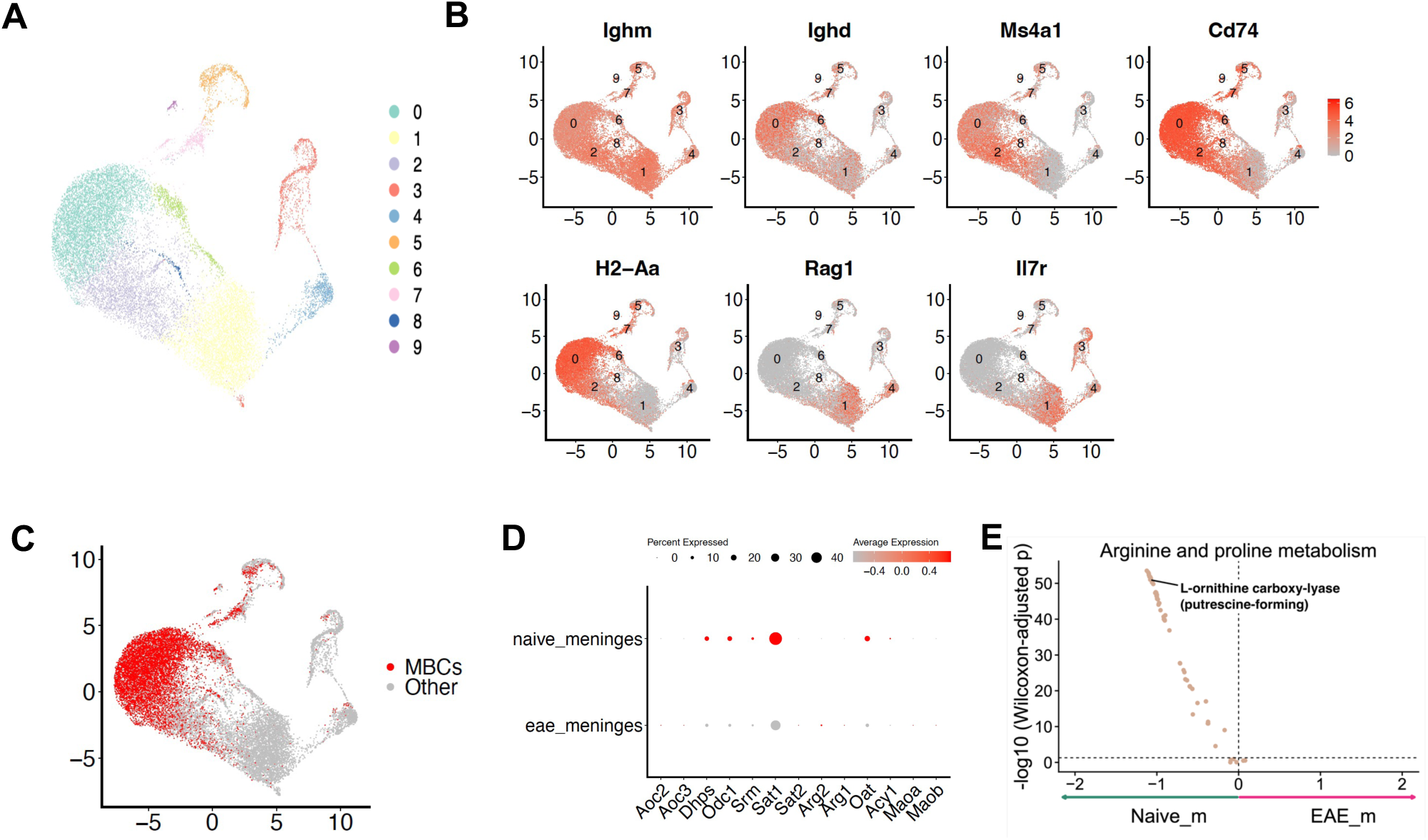
Polyamine metabolism is suppressed in activated meningeal B cells. Four C57BL/6 wildtype female and male mice were either immunized with MOG_35-55_/CFA to induce EAE or left untreated as naïve mice. At day 14, dura meninge were dissected and B cells (CD19+CD45+) were sorted by flow cytometry and subjected to scRNAseq (10x Genomics, **Methods**). **A)** UMAP. **B)** Expression of selected B cell related genes. **C)** Mature B cells (MBC) selected for further analysis based on the module score expression of *Ighd*, *Ms4a1*, *CD74*, and *H2-Aa* . **D)** Dot plot expression of genes in the polyamine/arginine pathway. **E)** Volcano plot showing Compass analysis comparing naïve and EAE dural B cells.

Next, we focused on mature B cells that co-expressed *Ighd*, *Ms4a1*, *CD74*, and *H2-Aa* (**Methods** and **Figure 4C**). Naïve mature meningeal B cells expressed *Odc1* and *Sat1*, both rate-limiting enzymes of the polyamine pathway, as well as affiliated enzymes such as *Dhps*, *Srm* and *Oat* (**Figure 4D**). Mature meningeal B cells from EAE mice, on the other hand, expressed lower levels of all 5 enzymes. A small fraction of those B cells expressed *Arg2* (**Figure 4D**). To determine whether polyamine reactions were significantly altered by EAE, we performed a Compass analysis (**Methods**). We found that EAE significantly reduced 53 metabolic reactions of the arginine/proline metabolism pathway (Wilcoxon-adjusted p < 0.0001 and Cohen’s d < -0.5, **Supplementary Table S3**, and **Figure 4E**). Notably, putrescine biosynthesis, which is catalyzed by ODC1, was among the top differential metabolic reactions identified (Wilcoxon-adjusted p = 1.54E-22, Cohen’s d = -1.1).

### ODC1 inhibition expands meningeal B cells

To evaluate whether the suppression of polyamine biosynthesis plays a functional role in meningeal B cell function, we treated mice from EAE onset (day 7) with 0.5% difluoromethylornithine (DFMO), a pharmacological inhibitor of ODC1, in drinking water. At day 14, dura meningeal CD4^+^ Teff cells were significantly reduced upon DFMO treatment (**Figure 5A**, upper left panel), a finding consistent with our previous report that DFMO restricts T cell differentiation.^30^ This reduction persisted at the chronic phase (day 54) in the leptomeninges (**Figure 5A**, lower left panel). No differences were observed in the number of regulatory T cells in either compartment or time point (**Figure 5A**, upper and lower middle panels). In contrast, DFMO treatment significantly increased the number of GC-like B cells (GL7^+^FAS^+^) in the dura meninges at day 14, consistent with the suppression of polyamine pathways in activated B cells (**Figure 5A**, upper right panel). This expansion did not persist at day 54 in the leptomeninges, likely due to the decline in CD4^+^ Teff cell support. Thus, unlike T cells, meningeal B cells suppress polyamine metabolism in EAE and the global ODC1 suppression can expand GC-like B cells in EAE.

**Figure 5.**
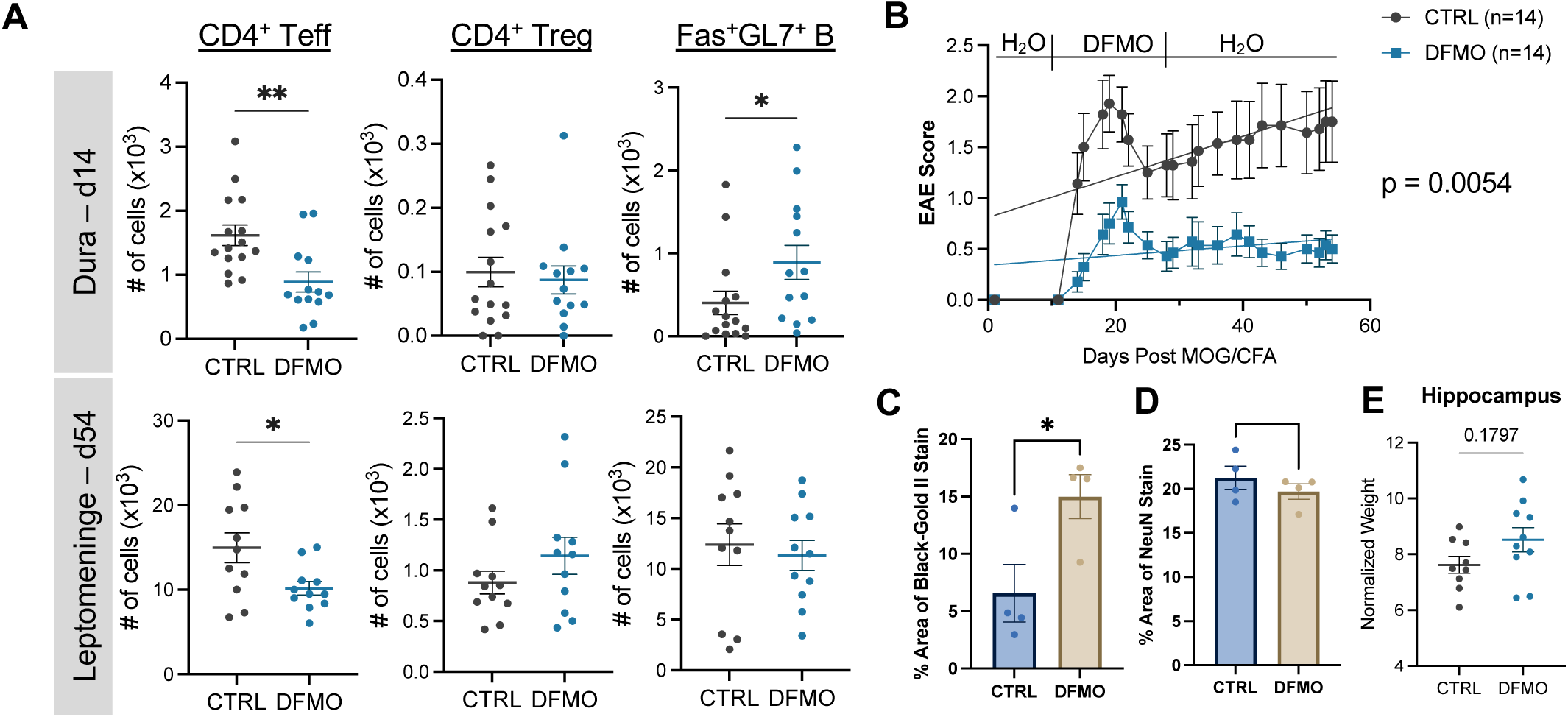
DFMO inhibits meningeal T cells but expands B cells in MOG_35-55_ EAE. C57BL/6 wildtype female and male mice were immunized with MOG_35-55_/CFA to induce EAE. DFMO were supplemented in drinking water (0.5%) from day 7-28. **A)** Meningeal cells were analyzed by flow cytometry at region and time as indicated. Male and female mice showed similar results and were pooled. **B)** EAE clinical score. **C-D)** Hippocampal myelination (C) and neurons (D) were quantified by immunohistochemistry. Each dot is one mouse and an average of 3 brain slices. **E)** Normalized hippocampus weight against the whole brain. Statistical analysis: A, C, D, and E used Student t test; B used linear regression analysis. * p < 0.05; ** p < 0.01.

Finally, consistent with MOG_35-55_ EAE being a T cell, and not B cell, driven model, therapeutic treatment of DFMO significantly reduced EAE severity (**Figure 5B**), reduced demyelination (**Figure 5C**) but had no effect on the number of hippocampal neurons or normalized weight (**Figure 5D, E**).

### B-cell intrinsic ODC1 restricts meningeal age-associated B cells and MOG_1-125_ EAE severity

To investigate the direct role of ODC1 on B cells, we first analyzed isolated dura meningeal B cells in culture with DFMO. DFMO promoted B cell proliferation when stimulated with LPS or IgM (**Figure 6A**). However, the effect of DFMO was negated when anti-CD40 was added in culture (**Figure 6A**), suggesting converging signals between CD40 signaling and ODC1 inhibition.

**Figure 6.**
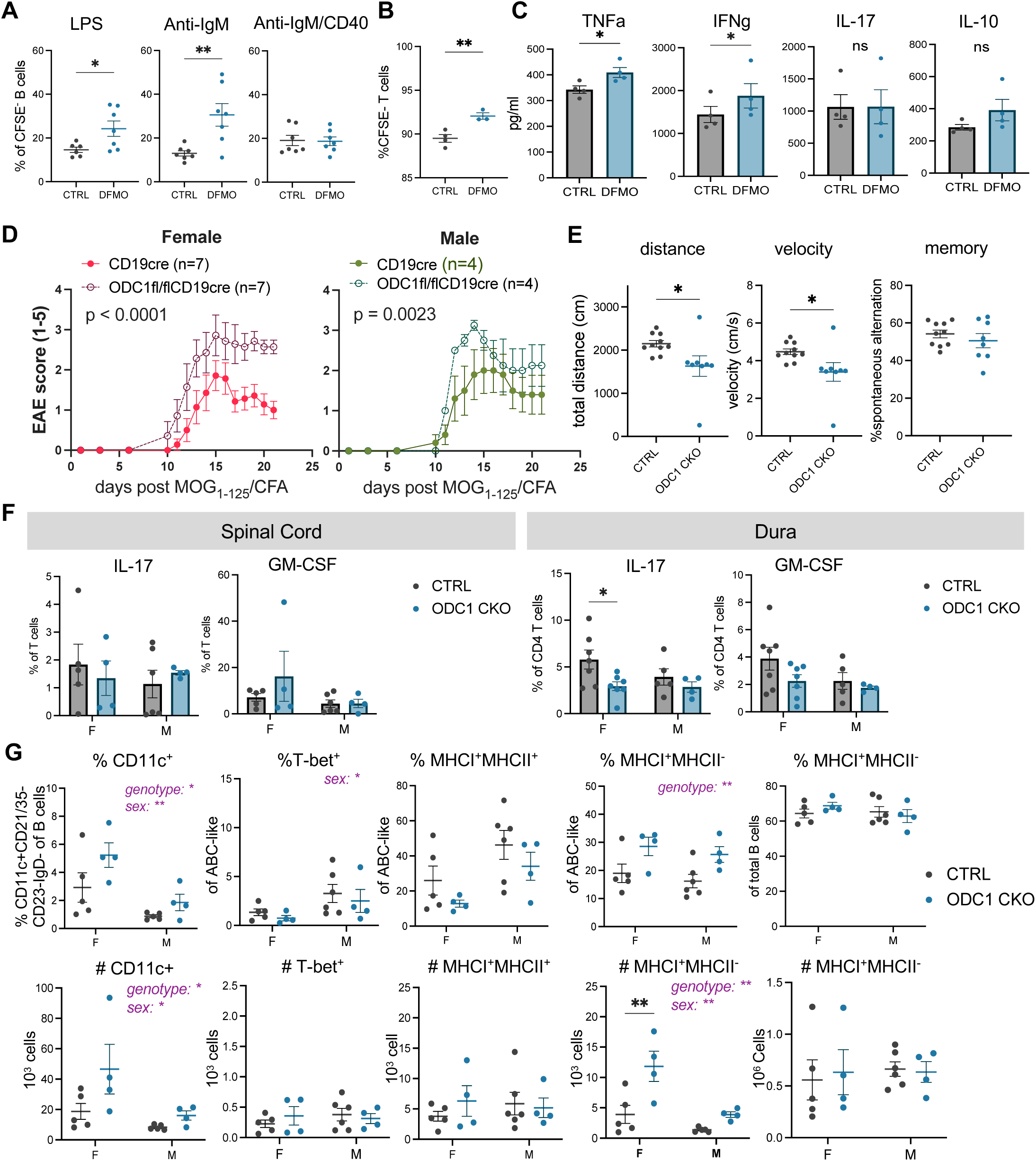
Intrinsic ODC1 restricts meningeal B cells function and MOG_1-125_ EAE severity. A-**C)** Dural meninges from naïve C57BL/6 female mice were dissected and B cells were enriched using magnetic beads and stimulated in vitro with control (CTRL) or DFMO (1mM) for 24 hours. A) B cells were analyzed by flow cytometry. B) B cell stimulated with anti-IgM were replated with fresh media and splenic CD4^+^ T cells isolated from C57BL/6 female mice for 3 days and analyzed by flow cytometry. C) Supernatant from B were analyzed for cytokine expression by Legendplex. **D-G)** Mice were immunized with MOG_1-125_/CFA to induce EAE. D) EAE clinical score; E) Y-maze analysis was performed on day 9 post immunization on both female and male mice which showed similar trend and results were pooled; F) T cell intracellular cytokine expression was evaluated; G) dural meningeal B cells was analyzed from day 14 post immunization by flow cytometry. In all panels, each dot is a mouse. Statistical analysis: A-C and E used unpaired student *t* test; D used linear regression analysis; F-G used two-way ANOVA with Tukey’s multiple comparisons test. * p < 0.05; ** p < 0.01.

Next, we asked whether DFMO-treated meningeal B cells had altered ability to prime CD4 T cells. CD4 T cells division frequencies were significantly increased despite their small effector size, when cultured with B cells pre-treated with DFMO (**Figure 6B**). We characterized cytokines in the supernatant of the B-T coculture and found increased levels of TNFa and IFNg, but not IL-10 or IL-17, when B cells were pre-treated with DFMO (**Figure 6C**).

To determine disease impact, we generated mice with a conditional deletion of *Odc1* in B cells (CD19^cre^ODC1^fl/fl^, ODC1 CKO). The MOG_1-125_ EAE model was selected because of its dependence on antigen presenting cells as opposed to MOG_35-55_ peptide immunization. We found that both female and male mice with ODC1 deficient B cells developed more severe EAE (**Figure 6D**). In a separate experiment, animal behavior was evaluated on day 9 prior to EAE score onset. ODC1 CKO mice were found to have significant reductions in the distance travelled and velocity when compared to littermate controls (CTRL, either CD19^cre^ or ODC1^fl/fl^) consistent with more severe EAE score that developed later (**Figure 6E**, left and middle panels). No difference was found in spontaneous alternation in a Y-maze test^37^ which evaluates spatial memory (**Figure 6E**, right panel).

To evaluate cellular changes, we isolated immune cells from the spinal cord and dural meninges. No differences were observed between the mice genotypes in IL-17^+^ or GM-CSF^+^ T cell frequencies in either tissue (**Figure 6F**). In contrast, there was an increased proportion and number of dural meningeal B cells with an age-associated-B-cell (ABC)-like phenotype: CD11c^+^CD21/35^-^ CD23^-^IgD^-^ at the onset of disease (**Figure 6G**). T-BET has been implicated in ABC biology in some^38^ but not all studies.^39^ We found no difference in the frequencies or number of ABC-like cells that express T-BET. Meningeal ABC-like cells from ODC1 CKO expressed higher frequencies of MHC class I single positive cells but not when co-expressed with MHC class II (**Figure 6G**, middle panels). The increase in MHC class I is not observed when gated on total B cells, suggesting a selective effect on ABC-like cells (**Figure 6G**, right panel).

## DISCUSSION

Meningeal inflammation is known to be associated with disease progression in MS, however a detailed understanding of the metabolic networks associated with this inflammation is lacking and presents an unmet need. This study identified peripheral metabolites related to meningeal inflammation and brain volume changes as defined by 7T brain MRI scans in a cohort of MS patients; and secondly evaluated the functional role of the polyamine pathway in regulating meningeal B cell function in EAE, a mouse model of MS. We found that 1) Polyamines were associated with both brain volume and meningeal inflammation in MS and EAE; 2) Polyamine metabolism was suppressed in activated B cells in EAE, in contrast to T cells that had increased polyamine activity^40^; 3) Pharmacological and genetic targeting of ODC1 resulted in an expansion of meningeal B cells particularly of the age-associated-like phenotype, which have been previously implicated in autoimmunity; 4) B cell intrinsic ODC1 limited EAE severity in the MOG_1-125_ EAE model.

The peripheral metabolic landscape was strongly associated with brain volume, especially of the hippocampus and caudate, alluding to their sensitivity and perhaps vulnerability to peripheral changes. Consistent with their general sensitivity, hippocampal volume loss is widely reported in healthy aging and central nervous system (CNS) diseases such as Alzheimer’s disease, Parkinson’s disease, and neuropsychiatric disorders. Although less studied, caudate volume decreases with age and is associated with memory decline and age-related neurodegeneration.^41,42^ The strong association of peripheral metabolites with brain regional volumes supports the relevance of metabolism, even peripheral metabolism, in CNS-specific disease. On the other hand, change in regional brain volume itself is not a marker of disease severity and typically alters brain connectivity in other parts of the brain that can be either adaptive or maladaptive in nature. Because these structures share anatomical proximity to the cerebral spinal fluid space, a better understanding of how hippocampal and caudate volume changes alter overall brain function will better inform the outside-in hypothesis of CNS health and disease.^43–45^

ODC1, the rate limiting enzyme of polyamine biosynthesis, plays a pleiotropic role in immune cell function and differentiation. We previously showed that ODC1 is critical for Th17 cell differentiation at the expense of regulatory T cell via epigenetic modifications.^30^ The differentiation of other CD4 helper T cell subsets is also dependent on polyamines and their role in the production of hypusine and histone acetylation.^46^ Consistent with a proinflammatory role in T cells, ODC1 inhibition was sufficient to alleviate MOG_35-55_ EAE, a T cell dependent model, when given either prophylactically,^30^ or therapeutically as shown in Figure 4B. In contrast, in macrophages, ODC1, as well as an upstream enzyme LACC1,^47^ restricts the development of an inflammatory state.^48^ The polyamine-dependent regulation of hypusination of translation factor eukaryotic initiation factor 5A promotes mitochondria function and selectively promotes alternatively activated macrophages.^49^ Consistent with this notion, ODC1 plays a protective role in macrophages in mucosal immunity against colitis and colon carcinogenesis.^50^ The role of ODC1 in B cells has not been studied previously. In the present study, we find that polyamine metabolism is suppressed in activated B cells. ODC1 plays a cell-intrinsic role in B cells as the perturbation of ODC1 expands meningeal B cells in the presence of either BCR signaling or LPS, but not when T cells help is present (anti-CD40). Since polyamines play a role in oxidative phosphorylation and IL-4-dependent pathways,^49^ we speculate that ODC1 is important in the initial metabolic reprograming of B cells during activation and germinal center formation, a process stabilized by anti-CD40 signals. Moreover, our data show that B cell intrinsic ODC1 restrains disease severity of EAE, in support of a protective role of polyamine metabolism in B cells and B cell-dependent autoimmunity model.

Age associated B cells (ABC) were first characterized by the Cancro and Marrack groups^51,52^ as splenic B cell subsets that accumulate with age and have since been found in other tissues including the CNS. ABC are heterogeneous and can be precursors to plasmablasts.^53^ ABC have unique cell surface markers (e.g. CD11b, CD11c^52^), transcription factors (e.g. T-BET^38^, ZEB2^54^) and are elevated in autoimmune and autoinflammatory diseases.^55^ CD11c^+^ ABC-like cells play a pathogenic role in a lupus model,^53^ and have also been shown to accumulate in EAE and Viral-EAE models of MS.^56^ The maintenance of the ABC population does not rely on B cell receptor (BCR) cross-linking alone, but instead depend on TLR7 and TLR9 signaling which can further synergize with BCR signaling.^55^ In addition to TLR7 and TLR9, cytokine signals such as IL-21 and IFNg can stabilize, whereas IL-4 can block, the emergence of ABC phenotype.^55^ Our study shows that B-cell specific deletion of ODC1 leads to an expansion of CD11c^+^ ABC-like cells in the meningeal compartment (Figure 5G). Interestingly the frequencies and numbers of of T-BET^+^ ABC were not different in the absence of ODC1 in B cells. Previous work has shown that while IFNg and TLR-signals can drive T-bet expression, CD11c expression is independent of T-BET but can be induced by IL-21.^57^ In fact, T-BET was found to be dispensable for ABC formation in some autoimmune diseases.^39^ Thus ODC1 restricts the development of ABC-like B cells in meninge. Whether ODC1 is connected to IL-21 signaling in B cells requires further investigation.

DFMO, an inhibitor of ODC1, is an FDA-approved drug for treating African sleeping sickness, and more recently, pediatric neuroblastoma. There is an increasing interest in the effect of DFMO in autoimmunity. Our work highlights a divergent role of the DFMO when targeting T and B cell responses. While ODC1 perturbation restricts effector CD4 T cells, it promotes the expansion of B cells, especially those with age-associated-like phenotype in the meninges during EAE. These findings suggest that DFMO may have a more pronounced effect in T cell-driven autoimmune responses, while its impact will be complicated in B cell-mediated pathology.

## STUDY LIMITATIONS AND FUTURE STUDIES

The present analysis of human samples is under powered for the evaluation of sex-, age- and treatment-specific effects which are important drivers of MS pathology. While ODC1 has divergent functions in T and B cells, the underlying mechanisms remain elusive and should be addressed in future studies.

## MATERIALS AND METHODS

### MRI Acquisition and analysis

MRI scans were acquired and analyzed as previously described (Zurawski et al. MSJ, Zurawski et al, JON, Callen et al JON). Briefly, a MAGNETOM Terra scanner (Siemens, Erlangen, Germany) equipped with a single-transmit/32-channel receive head coil was used for the brain 7T MRI, following a consistent scan protocol across subjects. This included T1-weighted (T1w) magnetization-prepared 2 rapid gradient-echo (MP2RAGE) and 3D T2-weighted FLAIR sequences, each with isotropic voxel dimensions of 0.7 × 0.7 × 0.7 mm^3^. Post-contrast imaging was conducted after an intravenous injection of 0.1 mmol/kg gadoterate meglumine with an approximate 10-minute delay between contrast administration and the start of FLAIR imaging. Experts who performed lesion identification and tracings, and the operator who performed automated analysis were blinded to clinical information.

A neurologist experienced in neuroimaging (JZ) manually compared pre- and post-contrast sagittal FLAIR images to detect leptomeningeal enhancement (LME). Scans were inspected for hyperintense signal on post-contrast FLAIR that were absent on pre-contrast FLAIR as previously published. Confirmed LME foci were counted. Cortical lesions (CL) were required to be hypointense on T1w-MP2RAGE compared to adjacent normal-appearing cortex, at least 1 mm wide, and distinct from blood vessels. The CL rater was blinded to LME status. FLAIR hyperintensity corresponding to T1w-MP2RAGE hypointensity often provided confirmation of CLs, though was not strictly necessary as FLAIR cortical ribbon artifacts can obscure lesions. CL volume was quantitated using a semi-automated edge-finding tool in Jim 7.0 software (Xinapse Systems Ltd, West Bergholt, UK; http://www.xinapse.com).

Whole-brain T2 hyperintense lesion volume (LV) was traced by a trained technician (YJ) using the Jim 7.0 software edge-finding tool on pre-contrast sagittal T2 FLAIR sequences.^10^

Automated MRI analysis was performed by an experienced computer scientist (RC). As detailed previously^58^, a customized pipeline was developed in-house for structural segmentation from 7T MRI scans. Initial “out of the box” DGM segmentations from FSL-FIRST (v5.0.9, Oxford, UK, https://fsl.fmrib.ox.ac.uk/fsl/fslwiki/FIRST) using the MP2RAGE images did not generate adequate DGM contours. Thus, we developed a pre-processing workflow involving noise suppression. We then applied the resulting UNI images from MP2RAGE reconstruction to the FSL-FIRST pipeline. Hippocampal volume segmentation was performed using FreeSurfer (v7.2.0, https://surfer.nmr.mgh.harvard.edu/fswiki). Pre-processed, denoised MP2RAGE images were used as input for hippocampal segmentation. Normalized brain parenchymal volume (BPV) and each subject’s whole brain/head normalization factor were obtained after applying the same pre-processing workflow using a fully automated segmentation pipeline (SIENAX v5.0, https://fsl.fmrib.ox.ac.uk/fsl/fslwiki/SIENA). The raw DGM and hippocampal volumes were multiplied by the normalization factor for each subject. Output segmentation files were reviewed for quality with any poor segmentation cases discarded as missing data.

### Metabolomics

#### Data acquisition

Human plasma samples were subjected to Liquid chromatography mass spectrometry at the Broad Institute (Cambridge, MA, USA)). Mouse brain tissue metabolomics for targeted measure of polyamines were performed at the Sickkids metabolomics core (Toronto, ON, Canada).

#### Metabolomics analysis

Downstream integration of metabolomic and MRI imaging data was performed in R version 3.4.4. Any variables with more than 30% missing values were excluded from analysis. Missing values for remaining continuous variables were imputed using the missForest package,^59^ and metabolite data was log-transformed with mean scaling. In order to examine the relationship between metabolite and brain pathology, we constructed a spare partial least squares (sPLS) regression model using the R mixOmics package.^60^ This method implements simultaneous dimension reduction and variable selection, allowing for robust associations to be made between a subset of metabolite predictors (X input matrix) and a set of outcome variables (Y outcome matrix). Input matrices were corrected for patient age using the *naiveBayesLM* function from the WGCNA package.^61^ LME foci number and another unbiased version using all imaging variables including regional brain volumes (hippocampus, deep gray matter, globus pallidus, thalamus, putamen, and caudate) and performing Y-variable regularization. The number of components for the model as well as the number of selected X variables was tuned using 5-fold, 5-repeat cross validation (CV), with optimal values chosen via the Q^2^ and maximum correlation criteria (measure = *“cor”*), respectively. sPLS model was constructed using the optimal number of components and variates in regression mode via the *spls* mixOmics function. Model stability was enforced via additional ensemble averaging of the optimal model outputs (loadings, coefficients, variate coordinates) across 50 total models constructed in this manner. Important variables were extracted from the ensemble model according to loading contribution via the *selectVar* function, and regression coefficients were obtained from the *predict.spls* function executed on the same model. Average Clustered Image Maps (CIMs) were generated using the *cim* function with the first 2 components of all unbiased sPLS model as input; each value in the heatmap (j, k) corresponds to the scalar product value between every pair of vectors in the first 2 components representing the variables X_j_ and Y_k_ on the axis defined by U_1_ and U_2_, where U_i_ is the i-th X variate. A cutoff of 0.2 was used for pairwise correlations, indicating at least one value per row must have an absolute value > 0.2 in the final output matrix. Network associations were calculated on metabolites in the first components of the ensemble sPLS model using bootstrapped Spearman correlation (999 permutations), with a hard threshold of |R|>0.3 and a corrected p-value < 0.05 applied to obtain the final network. Subsequent visualization of the metabolite network was performed using the igraph and qgraph packages.^62^

### Mice

Male and female wildtype C57BL/6 mice were bred in house and applied to experiments between 6-12 weeks of age. ODC1^fl/fl^ were gifted by Dr. Erika Pearce’s group and crossed to CD19^cre^ (both C57BL/6 background) and bred in house. Sex and age-matched littermate control mice (CD19^cre^ or ODC1^fl/fl^) are compared to ODC1^fl/fl^CD19^cre^ mice for all experiments. All mouse experiments followed guidelines outlined and animal user protocols approved by the Animal Care Committee at Sunnybrook Research Institute in Toronto, Ontario, Canada.

### EAE induction and DFMO treatment

MOG_35-55_/CFA or MOG_1-125_/CFA emulsion (Hooke’s lab) was injected at 100ug/mouse at both flanks subcutaneously to induce EAE followed by intraperitoneal injection of pertussis toxin (110ng/mouse) in 4 hours and 28 hours. Mice were monitored for clinical symptoms. DFMO (Selleck chemicals, 0.5% weight-gram/volume-100ml) was supplied in drinking water 7-30 days post immunization or as indicated.

### Y maze test

Behavioral test was performed in the morning in a specialized noise-, temperature-and light-controlled behavioral room. Mice were habituated in the room for at least an hour prior to the test. Each mouse was allowed to roam freely in a Y-maze chamber for 8 minutes following a published protocol.^37^ Mice movement was captured by an overhead camera and data was acquired and analyzed by Ethovision Software.

### Tissue/cell preparation

Dura, leptomeninges, and inguinal lymph nodes were collected after intracardiac perfusion with 10ml PBS. Meninge tissue was enzymatically digested on the shaker for 20min at 350rpm at 37℃ with Collagenase D (2.5 mg/ml) and DNase I (1mg/ml). Cells were filtered with 70µm strainers to obtain single cell suspension for downstream application. Inguinal lymph nodes were mashed directly on the 70µm strainers for cell isolation. Cells were either sorted by flow cytometry or enriched with magnetic microbeads for downstream applications.

### B cell culture

B cells were enriched from dural meninge using CD19 microbeads (Miltenyi Biotech). Enriched cells were seeded in 96-well round bottom tissue culture plate at a concentration of 2.5 × 10^5^/ml in complete media (RPMI1640, 10% heat inactivated Fetal Bovine Serum, NEAA, Glutamate, Pyruvate, beta-mecaptoethanol) with anti-IgM (Jackson ImmunoResearch, 10ug/ml), anti-CD40 (Jackson ImmunoResearch, 1ug/ml), LPS (Sigma, 100ng/ml) or combinations. For proliferation assay, CFSE (Thermofisher, 0.5uM) was used to label enriched B cells followed by 3 days of culture and flow cytometry analysis. For B-T coculture, B cells were cultured with anti-IgM and either control or 1mM DFMO for 24 hours. Cells were washed and replated with CFSE labeled enriched CD4 T cells (Miltenyi Biotech) in fresh media for 3 days. Co-cultured cells were analyzed by flow cytometry, and culture supernatant was analyzed by legendplex (Biolegend).

### Antibodies

Staining of cell surface markers was performed using Brilliant Stain Buffer (BD Biosciences) to minimize fluorochrome interactions. Cells were incubated for 30 minutes at 4°C in the dark with a panel of fluorochrome-conjugated monoclonal antibodies, all obtained from BD Biosciences unless otherwise specified. The following antibodies and dilutions were used: fixable viability dye 510 (1:1000) for live/dead discrimination, anti-CD45 FITC (1:1000), anti-CD19 RB545 (1:400), anti-B220 BUV496 (1:200), anti-CD2 BV650 (1:400), anti-CD11b RY605 (1:400), anti-CD8a BV605 (1:400), anti-CD11c RB780 (1:100), anti-CD4 PerCP (1:200), anti-CD3 APC (1:200), anti-CD21/CD35 BUV496 (1:500), anti-CD40 BUV395 (1:200), anti-CXCR4 BUV805 (1:200), anti-CXCR5 RY610 (1:200), anti-IgA BV605 (1:200), anti-IgM RB613 (1:200), anti-IgD BV711 (1:400), anti-T-bet RB705 (1:50), anti-IA-IE APC/Fire 810 (Biolegend 1:400), and anti-MHC I H2Db PE-Cy7 (Thermofischer 1:400). Following surface staining, cells were fixed overnight using a commercially available fixation buffer. The next day, cells were permeabilized using a standard permeabilization buffer (BD Biosciences) and subsequently stained for intracellular markers, including T-bet. After intracellular staining, cells were washed and resuspended in staining buffer prior to acquisition. Data were collected using a spectral flow cytometer, and appropriate compensation and fluorescence-minus-one (FMO) controls were included for analysis.

### scRNAseq

#### Sample preparation

Single-cell libraries were generated using the Chromium Next GEM Single Cell 5’ Reagent Kit v2 with Feature Barcode technology for Cell Surface Protein (10x Genomics, PN-1000263), following the manufacturer’s protocol. Single-cell suspensions were obtained from four individual mice (2 males and 2 females) per experimental group and labeled with unique sample-specific barcoded antibodies (hashtag oligonucleotides) to enable multiplexing. Equal numbers of labeled cells from each mouse were pooled at a 1:1:1:1 ratio to a final concentration of 1 × 10⁶ cells/mL. A total of 16,500 cells were loaded into one lane of a Chromium GEM Chip K, targeting an expected recovery of approximately 10,000 cells. Gel Bead-in-Emulsions (GEMs) were generated using the Chromium Controller (Wang Lab, Biological Sciences Platform of Sunnybrook Research Institute), followed by reverse transcription and cDNA amplification. For each reaction, 50 ng of amplified cDNA was used for library construction according to the manufacturer’s instructions. Equal molar amounts of gene expression and cell surface protein libraries were pooled and sequenced on the Illumina NovaSeq X 10B flow cell platform (The Centre for Applied Genomics, The Hospital for Sick Children, Toronto).

#### scRNA-seq analysis

Demultiplexing, alignment, barcode extraction, and UMI-Collapsing were performed by Cell Ranger Pipeline (v7.2.0) from 10X genomics. QC, data normalization (NormalizedData) and scaling (ScaleData), dimensionality reduction (PCA and UMAP), clustering (FindClusters), and differential gene expression (FindAllMarkers and FindMarkers) were performed with the Seurat R package [v4.4.0].^63^

We removed low-quality cells and cell doublets determined by abnormal unique number of genes (>2500 or <200) and high mitochondria content (>10%). Results were cleaned up further by selecting cells expressing *Cd19, Cd79a,* and *Cd79b* for downstream analysis (Broschi et al., 2021; Diddens et al, 2024; Schafflick et al, 2021). Data were normalized with the ‘LogNormalization’ method, and highly variable genes were computed by the function FindVariableFeatures in the default setting using method ‘svt’ and nFeatures equals to 2000. Those highly variable genes are used to perform PCA after scaling of the data. Jackstraw analysis and Elbow plot were used to select distinguished principal components, and the top 10 PCs were included for downstream analysis. Samples were then integrated using the Harmony algorithm.^64^ The cells were clustered via FindNeighbors and FindClusters functions, setting resolution to 0.3 using the reduction result from harmony. UMAP was used to visualize the results.

### Cellular Metabolic Prediction Analysis

The Compass [v1.0.0] (Wagner et al., 2021) algorithm was used to predict cellular metabolic activities within mature meningeal B cells (MBCs). Gene expressions were normalized, and 500 MBCs of each sample were selected to run Compass using the metabolic model ‘Mouse1’. MBCs are B cells that have a signature score greater than 1.5 calculated by average expression of *Ighd, Cd74, H2-Aa.*^14,65,66^ Post-compass analysis was based on the documentation provided by Wagner Lab, including normalization, scaling, and an unpaired Wilcoxon rank-sum test. Python packages NumPy [v2.1.1], Pandas [v2.2.3], and Matplotlib [v3.10.1] were used for data analysis and visualization.

## ACKNOWLEDGEMENTS

This work was supported by grants from the National Multiple Sclerosis Society (Career Transition Fellow Grant, Faculty Phase, TA-2101-37225) and Canadian Institute of Health Research (Project Grant #498150) awarded to CW.

## AUTHOR CONTRIBUTIONS

CW, JZ, and RB conceived, designed and supervised the study. CW wrote the manuscript with significant help from JZ and AT and contributions from all co-authors. JZ performed and analyzed the 7-MRI human data and oversaw patient recruitment along with RB. MP, AT, CW designed, conducted and analyzed mouse experiments with help from SP. JW analyzed the mouse single cell RNAseq data. KC performed scRNAseq sample preparation and acquisition. CC performed/supervised the human plasma metabolomics LC/MS acquisition and post-processing analysis. XX performed initial statistical analysis of human metabolomics data with AK’s supervision. MG performed the final statistical analysis of human metabolomics data with CW’s supervision.

